# Fluid-Squid: DIY Multiplexed Imaging of Cells and Tissues

**DOI:** 10.1101/2025.10.09.680291

**Authors:** John W. Hickey, Hongquan Li, Chiara Caraccio, You Yan, Kevin Marx, Peter Martin, Hope T Leng, John Fayiah, Elvis Towalid, Bonnie Dighero-Kemp, Garry P. Nolan, Manu Prakash, David R. McIlwain

## Abstract

Recent advances in multiplexed single-cell characterization have revolutionized our insight into cell biology, but many available technologies remain limited by high costs or a lack of customizability. To address these challenges, we developed Fluid-Squid, a cost-effective, quantitative imaging platform that integrates automated fluidics to support customizable, do-it-yourself multiplexed imaging workflows. Using Fluid-Squid, we successfully imaged fresh frozen human intestinal tissues with a 36-plex oligonucleotide-barcoded antibody panel and further demonstrated the feasibility of lyophilizing such multiplexed panels. We also adapted existing multiplexed imaging workflows to characterize individual cells to identify immune cell populations, phenotype, and antigen-specific cells from mouse splenocytes and human peripheral blood mononuclear cells (PBMCs) with a 39-antibody panel. To further expand its utility, we developed a barcoding strategy that allows for the pooling and simultaneous staining of multiple samples, reducing time, costs, and batch effects in single-cell experiments. This approach facilitated rapid titration to optimize antibody concentrations and assess the impact of various blood preparation methods on cell type retention. Overall, our work provides a new open-source framework for automated fluidics and microscopy in a flexible, cost-effective platform, empowering adaptable multiplexed characterization of both single cells and tissues.

## Introduction

The past decade has seen remarkable advancements in single-cell biology, driven in part by the development of multiplexed imaging technologies. In particular, spatial-omics approaches—both spatial proteomic (*1*) and transcriptomic (*2*) methods—have driven insights into cellular function and organization within tissues for (*3*), cancer (*4*), and various diseases (*5, 6*). These technologies enable simultaneous measurement of 50 or more markers at single-cell resolution. Consequently, single cell and spatial-omics techniques have been adopted by large-scale national and international tissue mapping consortia and have led to numerous commercialized systems.

However, despite their commercialization, many of these systems remain prohibitively expensive and complex, limiting their accessibility or adaptability to modification in protocols. For instance, most of these platforms rely on automated fluidics for reagent exchange during multiplexing protocols (*7–10*) but do not lend well to adaptations to new procedures. While such automation increases throughput and reproducibility, it also adds substantial cost and reduces flexibility, as many commercial systems are designed to work with proprietary reagents and protocols. The lack of customizability in these systems constrains their use for researchers who require bespoke experimental setups, need to integrate new reagents, or want to establish new techniques.

In addition to spatial technologies, single-cell suspension-based approaches such as mass cytometry (CyTOF) (*11*) and spectral flow cytometry (*12*) have emerged as powerful tools for profiling large numbers of markers (40+) on dissociated cells. However, like spatial-omics platforms, these methods require access to large, complex, and costly instrumentation, limiting their deployment to well-equipped labs or specialized core facilities. This reliance on expensive infrastructure presents a significant barrier, particularly for researchers conducting fieldwork in resource-limited settings, such as infectious disease outbreak studies such as Ebola that require the analysis of peripheral blood mononuclear cells (PBMCs) in situ (*13*).

Given these challenges, there is a pressing need for accessible and customizable solutions that democratize multiplexed imaging technology. In response to this need, we developed Fluid-Squid, an open-source platform for multiplexed imaging that integrates automated fluidics with low-cost high-performance microscopy systems. Fluid-Squid is designed to empower researchers to conduct high-parameter imaging experiments in both tissue and single-cell contexts. With flexible open-source software control, Fluid-Squid enables do-it-yourself (DIY) multiplexed workflows offering researchers the flexibility to customize their experimental protocols, incorporate novel reagents, and perform cost-effective assays without the need for expensive large-footprint equipment.

We demonstrate the versatility of Fluid-Squid through its application in characterizing tissue sections and immune cell populations, phenotypes, and antigen-specific cells in dissociated mouse splenocytes and human PBMCs. Additionally, we present a barcoding strategy that allows for the pooling and simultaneous staining of multiple samples, significantly reducing time, costs, and batch effects in single-cell experiments. This open-source platform bridges the gap between high-cost commercial systems and the need for customizable, scalable multiplexed imaging approaches, making advanced single-cell and tissue-level characterization rapidly accessible to a wider scientific community.

## Results

### Fluid-Squid: Low-cost, open-source platform for multiplexed imaging

We recently developed Squid (Simplifying QUantitative Imaging platform Development and deployment) (*14*). This platform is a cost-effective, quantitative imaging platform featuring a comprehensive set of hardware and software for advanced widefield microscopy techniques like flat-field fluorescence excitation microscopy. Its open, modular design not only simplifies customization and programming for diverse experiments but also enhances accessibility, making advanced microscopy and deep learning-based methods more feasible for point-of-care or low-resource settings. Squid has now been field tested in more than a dozen applications including confocal microscopy, expansion microscopy and zero-gravity imaging (*14, 15*).

Building on this platform, we created Fluid-Squid, which integrates automated fluid handling (**Fig. 1A**) with automated imaging to enable multiplexed imaging. This setup is compact (**Fig. 1B**), requires minimal lab bench space, and includes dedicated software to control fluid flow from multiple ports (**Fig. 1C**). It is robust, portable, and has been validated through transport and deployment in partner institutions from the United States to Liberia where it was used (**Fig. 1D)**. A photo of a newer generation of the fluidics system coupled with a Squid spinning disk confocal is also shown in **Fig. 1D**. Overall, Fluid-Squid allows large-scale, multiplexed experiments, such as antibody-based imaging, to be performed autonomously with both fluidics and imaging fully integrated.

**Figure 1.**
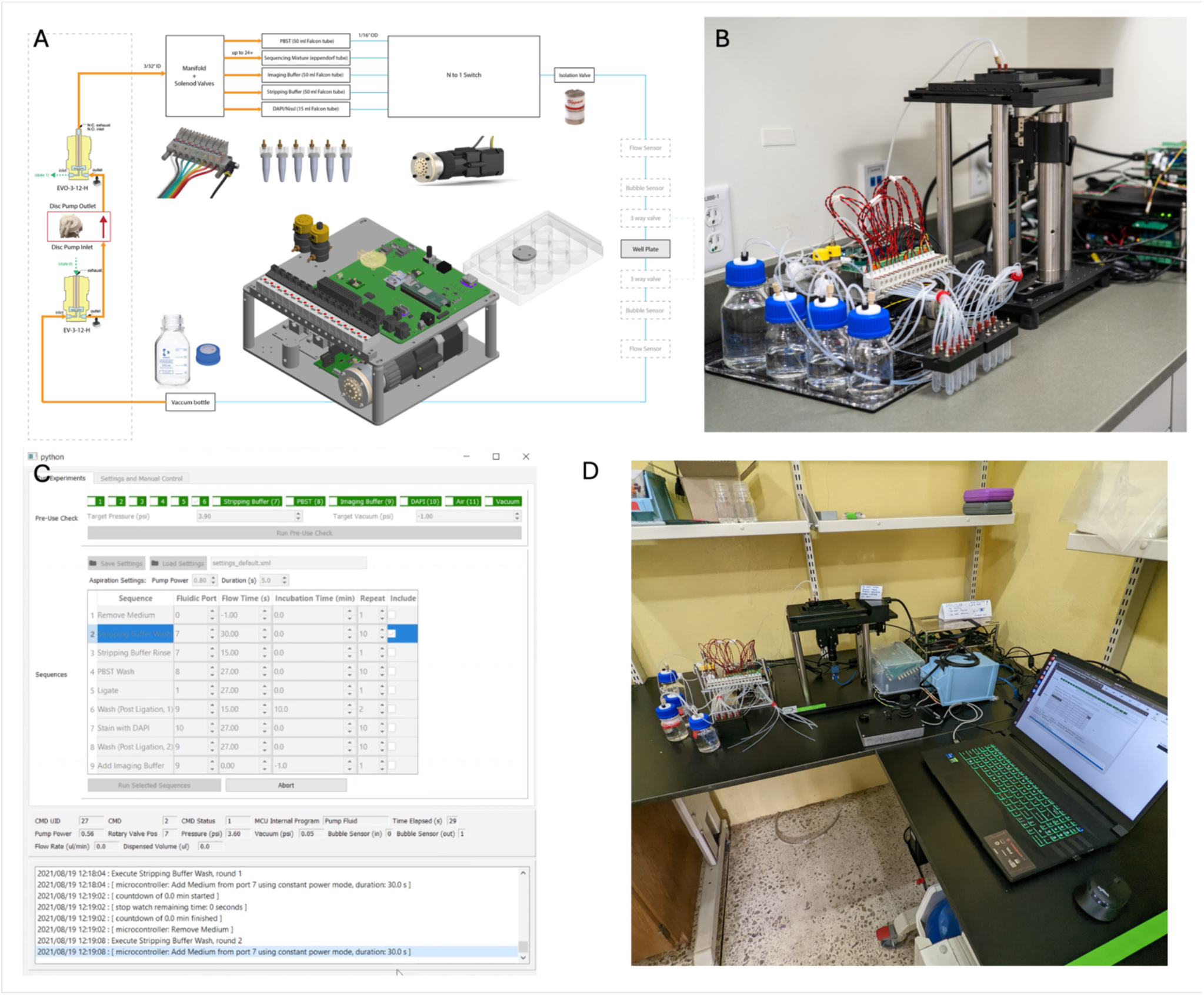
Fluid-Squid a customizable, open setup which integrates automated fluid handling with automated imaging to enable multiplexed imaging. A) Schematic and core components CAD of the generation 1 fluidics system, B) Photo of the fluidics system coupled to a Squid inverted microscope, C) Graphical user interface of the fluidics system, D) Photo from trip to Liberia; the same instrument from Figure B set up at Liberia Institute for Biomedical Research; a newer generation of the fluidics system coupled to a Squid spinning disk confocal.

The first-generation Squid fluidics system makes use of pressure driven flow to deliver reagents from sealed reservoirs to sample chambers like well plates. The pressure (up to 5 psi) is generated by an on-board piezo disc pump (Ventus). The same pump is also used to generate vacuum for aspiration. Pressure sensors are used for closed loop pressure control and for monitoring purposes (e.g. ensure reservoirs are properly sealed during automated pre-use check). A media isolation valve is used in the delivery path to ensure flow only starts after set pressure is reached for delivery of repeatable volumes. Flow sensors can be connected for additional monitoring, closed-loop control, and calibration purposes. Reservoirs for cycle-specific reagents are DNA LoBind eppendorf tubes connected to a custom machined manifold with o-rings and threaded adapters to lock the tubes in place.

In a second generation system, we used a design similar to the ones reported in (*16, 17*), where a syringe pump is used to pull reagents through a closed flow cell or aspirate reagents and dispense to open chambers. In this case the reagents can be stored in open instead of sealed reservoirs like well plates. The controller for the second-generation system also includes interfaces for multiple selector valves for more expandability (e.g., adding a selector valve for delivering reagents to multiple sample chambers).

The Squid microscope is constructed as described previously in (*14*) except that 1) a motorized stage for well plate with 80 x 120 mm travel is used for well plates, 2) Squid 850 nm laser autofocus is integrated, 3) the 4 color laser engine is further optimized., and 4) instead of using the more expensive 561 nm line, a 530 nm diode laser is used for cy3. The total BOM (bill of materials) cost of the microscope and the fluidic system put together between 2021 and 2022 is within $30k.

### Multiplexed Imaging of tissues using Fluid-Squid

Multiplexed imaging has transformed our understanding of the organization of tissues by enabling simultaneous profiling of dozens of markers at a single time (*3*). Computational tools that extract molecular information encoded as fluorescent signals, have enabled insight into cellular interactions and multicellular organization of tissues (*18–24*). We have previously developed CODEX (Co-Detection by indEXing) multiplexed imaging to profile cellular organization of tissue sections (*10, 25–27*). This technique leverages oligonucleotide-barcoded antibodies and cycling on and off fluorophore-conjugated, complementary oligonucleotides to iteratively reveal and image dozens of antibodies within tissue sections.

For the first time, we leveraged Fluid-Squid to perform CODEX multiplexed imaging with 36 DNA-barcoded antibodies within healthy human intestine tissue (*18*). We stained, imaged, and processed an area of the intestine of approximately 4.5 mm by 4.5 mm (**Fig. 2A**). The composite image of a subset of markers demonstrates consistent, high-quality staining across the entire section. Alpha smooth muscle actin effectively marked smooth muscle cells in both the muscularis externa and muscularis mucosa, cytokeratin identified epithelial cells in the mucosa, and CD56 highlighted nerve cells dispersed throughout the intestinal layers (**Fig. 2A**). Further analysis of the mucosal region revealed CD138 marking plasma cells, CD8 identifying CD8+ T cells, and Ki67 highlighting proliferating immune and epithelial cells (**Fig. 2B**). Similarly, other markers visualized aligned with known cell types and localizations within the intestine (**Fig. 2C-E**). These data indicate that Fluid-Squid is able to provide high-fidelity fluidics handling of cyclic fluid exchanges and integration with imaging of stained tissue sections required for CODEX.

**Figure 2.**
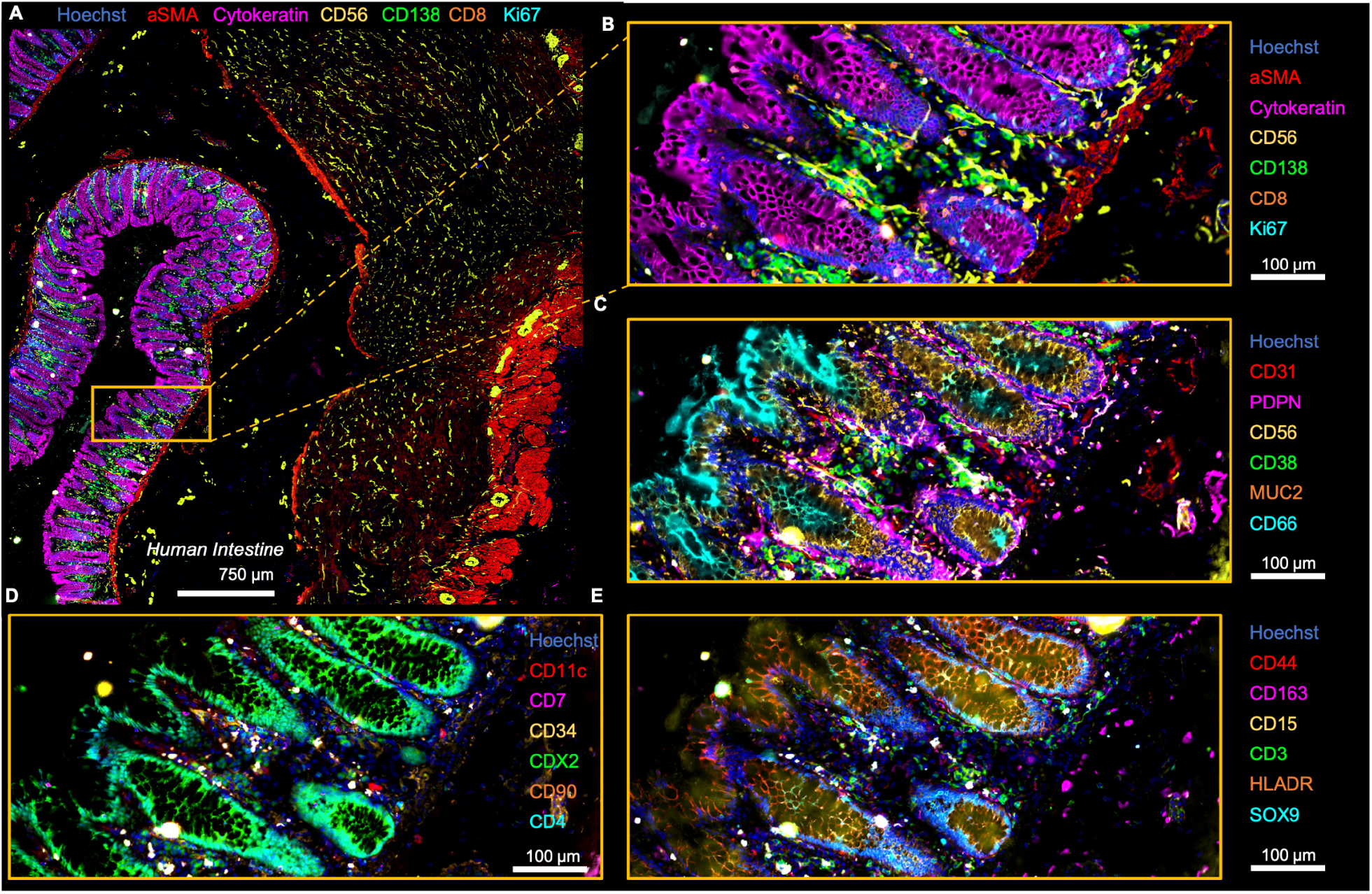
We used Fluid-Squid to perform CODEX multiplexed imaging of human intestine tissue. **A)** Imaging of a several mm^2^ area of tissue with single-cell resolution. **B-E)** magnified regions showing distinct marker sets from the same experiment, 6 of 36 markers shown per panel.

One key advantage of the Fluid-Squid system is its portability and flexibility. However, challenges arise in transporting and storing multiplexed imaging panels, particularly due to cold-chain storage requirements. To address this, we explored whether CODEX antibodies could be lyophilized and later reconstituted for staining. We tested this by comparing the performance of oligonucleotide-conjugated CD8 and B220 antibodies on mouse spleen sections: one group was used fresh, while the other was lyophilized and then reconstituted. The reconstituted antibodies performed comparably to those maintained in solution (**Fig. S1**). This suggests that lyophilization enables CODEX antibody panels to be portable and standardized for use across different staining protocols.

### Application of CODEX multiplexed imaging to dissociated single cells

We and others have mostly focused on using multiplexed imaging methods to analyze tissue samples (*4, 19, 20*). For example, our imaging of the human intestine or response to anti-cancer T cell therapies illustrates the powerful insights that microscopy-based approaches provide in understanding tissue architecture (*18, 19*).

Multiplexed analysis of single cells such as peripheral blood mononuclear cells (PBMCs) with techniques such as CyTOF (*11*) have also transformed our views into human immunology with measurement of the diversity, function, and composition of cells that relate to healthy and disease states. However, current techniques have several limitations. First, complex methods such as CyTOF or multispectral flow cytometry rely on expensive, large, elaborate setups. Second, other approaches such as single-cell RNA sequencing remain very expensive on a per-cell basis due to sequencing costs. Third, many of these multiplexed techniques lose important features such as cellular morphology or shape.

To address these limitations, we developed a technique that applies CODEX multiplexed imaging to dissociated single-cell populations. Our approach follows the general workflow of traditional multiplexed techniques like CyTOF, where individual cell populations are isolated and stained with a cocktail of tagged antibodies. However, instead of analyzing the cells through microfluidics, they are fixed onto a glass surface and imaged using CODEX cyclic multiplexed imaging (**Fig. 3A**).

**Figure 3.**
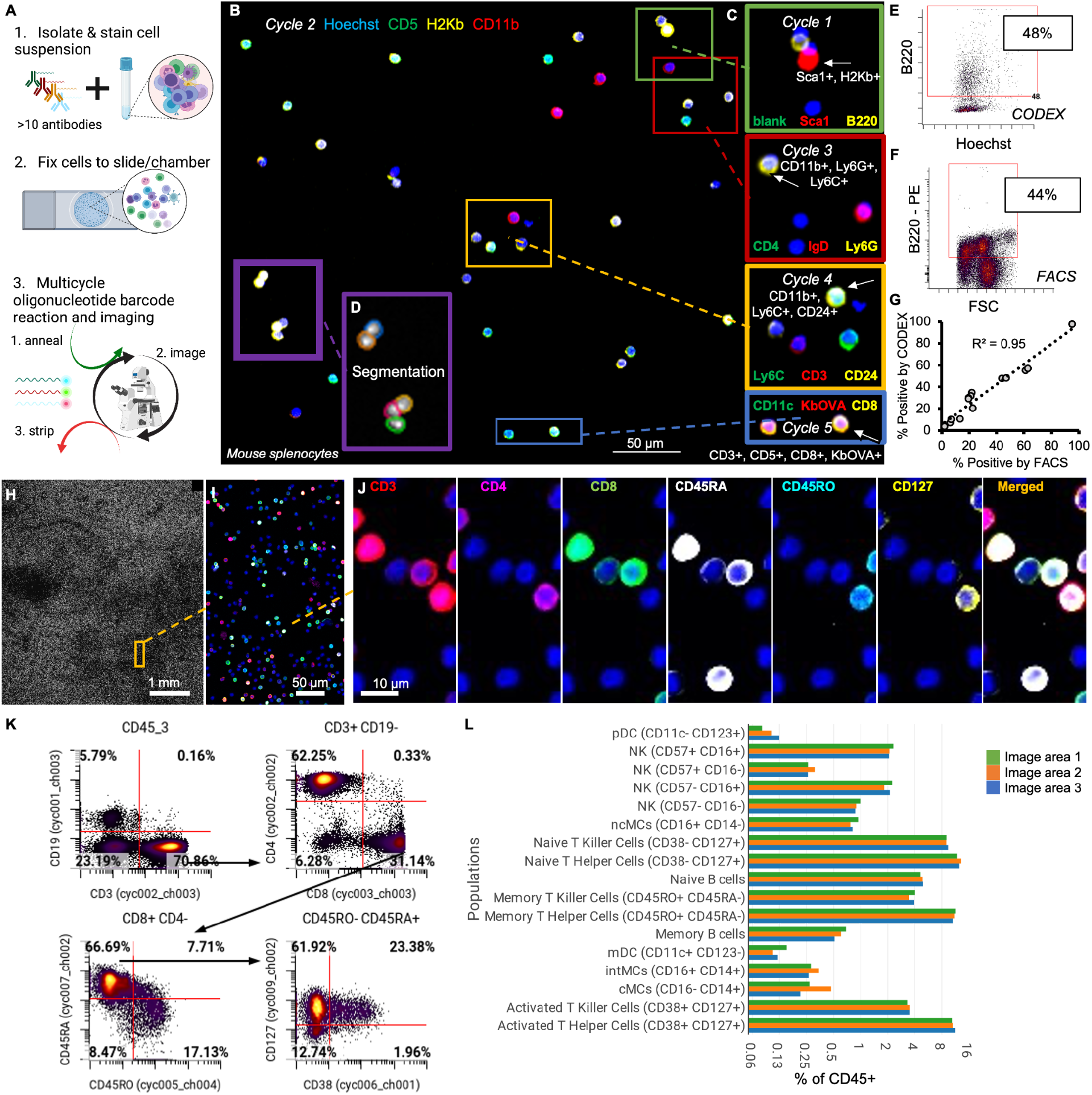
We adapted CODEX multiplexed imaging to profile dissociated single cells like splenocytes or PBMCs. **A)** Our method works by staining cells in suspension with a cocktail of oligonucleotide-barcoded antibodies and fixing stained cells to slides or staining chambers and then performing cyclic imaging by adding and removing complementary, fluorophore-tagged oligonucleotides. **B)** Representative region from imaging mouse splenocytes across five cycles of imaging with only 3 of the 14 markers shown and **C)** in magnified regions with additional markers shown across all five cycles with **D)** cell segmentation masks. **E-F)** Using the same samples, we compared the **E)** percent cells positive for B220 from our CODEX profiling compared to **F)** splenocytes positive for B220 using flow cytometry. **G)** Correlation of data derived from either CODEX or flow cytometry across a number of markers and cell populations. **H-L)** Profiling of millions of human PBMCs across **H)** entire slides that **I-J)** when magnified and overlaid with markers can demonstrate different subpopulations of T cells with 6 of 36 markers shown. **K)** Quantification and gating of imaging data to identify specific cell type populations that are **L)** consistent across multiple different regions of the image.

We first optimized and validated this technique using mouse splenocytes, applying a 14-antibody panel to cells in a 12-well glass plate. Manual cyclic fluidics were performed, and imaging was carried out with a commercial Keyence microscope (prior to adapting to Fluid-Squid as described in the following section). We efficiently captured fluorescent signals corresponding to distinct cell populations (**Fig. 3B**). We detect B cells (cycle 1), myeloid cells (cycle 2), neutrophils (cycle 3), T cells (cycle 4), and antigen-specific T cells (cycle 5) (**Fig. 3C**).

By performing cell segmentation (**Fig. 3D**) we quantified CODEX fluorescent signals at the single-cell level and compared them with flow cytometry results. To accomplish this, the same splenocyte sample was divided, stained with fluorophore-labeled antibodies, and analyzed using flow cytometry. Within our CODEX imaging data, approximately 48% of cells were labeled as B220+ (**Fig. 3E**), closely matching the 44% identified by flow cytometry with PE-labeled anti-B220 (**Fig. 3F**). This analysis extended to 14 additional proteins shared between both assays, yielding a strong correlation (r^2^ = 0.95) between populations identified by CODEX multiplexed imaging and flow cytometry (**Fig. 3G**).

We further applied this method to PBMCs, placing the cells on a slide compatible with existing automated CODEX fluidics and imaging protocols designed for analyzing tissues. This also enabled imaging of large areas (~50 mm^2^) of densely packed cells, resulting in data from hundreds of thousands of cells (**Fig. 3H**). We used a panel of 36 DNA-barcoded antibodies to delineate cell types and phenotypes, including distinct T cell phenotypes observed both in imaging data (**Fig. 3I-J**) and segmented single-cell data (**Fig. 3K**). A clear separation was evident between distinct cell type populations using markers like CD19 (B cells), CD3 (T cells), CD4 (CD4+ T cells), and CD8 (CD8+ T cells) (**Fig. 3K**). Continuous changes in protein expression, such as the decrease in CD45RA and simultaneous increase in CD45RO, were consistent with trends observed in other single-cell techniques like CyTOF (*27*) (**Fig. 3K**).

Finally, we analyzed spatial consistency by dividing the image into three areas and applying the same gating strategy (**Fig. S2**). Robust and consistent percentages of cell types were identified across different regions, indicating that our method reliably captures cell population information without introducing spatial batch effects (**Fig. 3L**). In total this represents a viable method for multiplexed characterization of dissociated single cells.

### Fluid-Squid for multiplexed imaging of single cells

Having validated the CODEX approach on both mouse and human dissociated single cells, we proceeded to adapt the assay to the Fluid-Squid platform. First, we confirmed that the assay could be successfully performed with manual buffer exchanges, solely performing imaging using the Squid setup. The image quality obtained from Squid was comparable to that from a commercial fluorescent Keyence microscope (**Fig. S3A**). We reserved a set of PBMCs from the same donor and analyzed them with flow cytometry. Quantitatively, we observed similar proportions of CD3+ cells from Keyence or Squid images as those that were stained with alexa-647 anti-CD3, analyzed via flow cytometry (**Fig. S3B**).

Next, we applied a full-scale 29-antibody panel to stain healthy human PBMCs and performed CODEX assay buffer exchanges and imaging in an automated manner using Fluid-Squid. Robust signals were observed from various cell populations within a subset of the total imaging area (**Fig. 4A**). We then performed cell segmentation across the entire imaging area, quantified the fluorescent signals, and conducted unsupervised clustering (**Fig. 4B**). This analysis identified 12 distinct cell types and phenotypes (**Fig. 4B-C**), associated with expected enrichment for cell type markers (**Fig. 4D**). By phenotyping in this manner, we were able to isolate individual populations, such as memory CD4+ T, B, and CD16+ NK cells, and examine specific marker distributions per cell visually. This allowed us to observe cell morphology, the overall intensity of each marker per cell, spatial distribution, cell polarization, and heterogeneity in cell type markers within the population (**Fig. 4E**).

**Figure 4.**
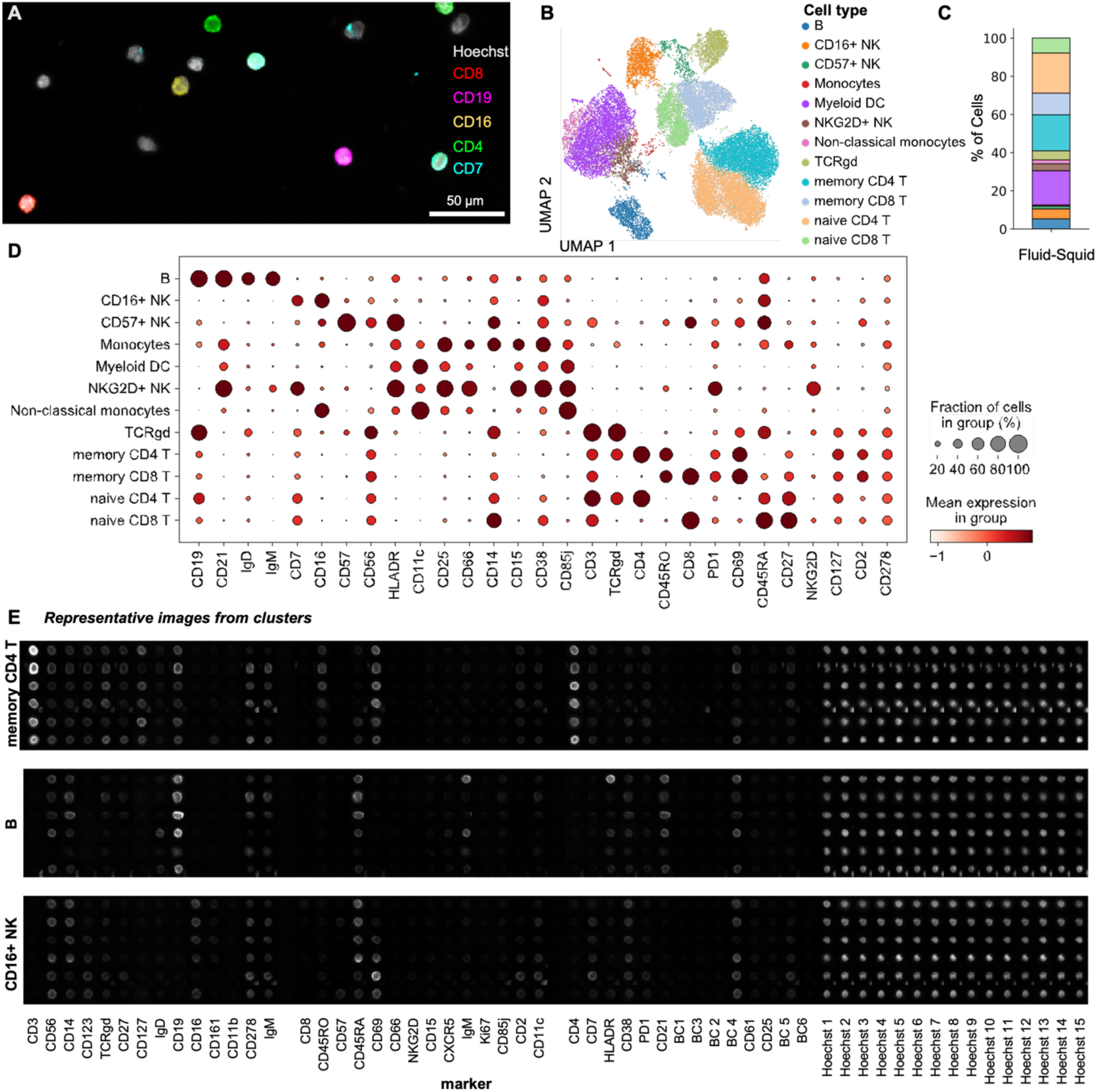
We used our Fluid-Squid set up to profile human PBMCs using a panel of 29 antibodies. **A)** A representative area of PBMCs containing various cell populations (image overlay for 5 of 29 markers used). **B)** UMAP plot of the cell types identified from multiplexed imaging of PBMCs, **C)** quantification within the sample, and **D)** dotplot highlighting marker expression on annotated cell types. **E)** Example of individual cell images for a few cells from the Memory CD4+ T cell population across all individual fluorescent channels used (including all the nuclear channels).

### Multiplexing sample analysis for large-scale single-cell studies

A major advantage of profiling PBMCs with multiplexed imaging is the ability to characterize large numbers of cells simultaneously. Many studies that examine immune responses in humans using PBMCs will enlist large cohorts of donors. Because costs scale with the number of assays run, these studies can become costly. To address this issue, we aimed to develop a sample barcoding strategy that would allow pooling cells and samples for simultaneous staining and imaging, thereby reducing costs, time, and batch effects.

We developed a barcoding strategy leveraging anti-CD45, a pan-immune cell marker (**Fig. 5A**), similar to those used in applications like CyTOF (*28*). To do this, we conjugated unique oligonucleotide barcodes to a distinct anti-CD45 antibody clone, generating unique set of anti-CD45 barcoding reagents. Individual cell populations were then stained with a unique anti-CD45 oligonucleotide combination before pooling for downstream staining.

**Figure 5.**
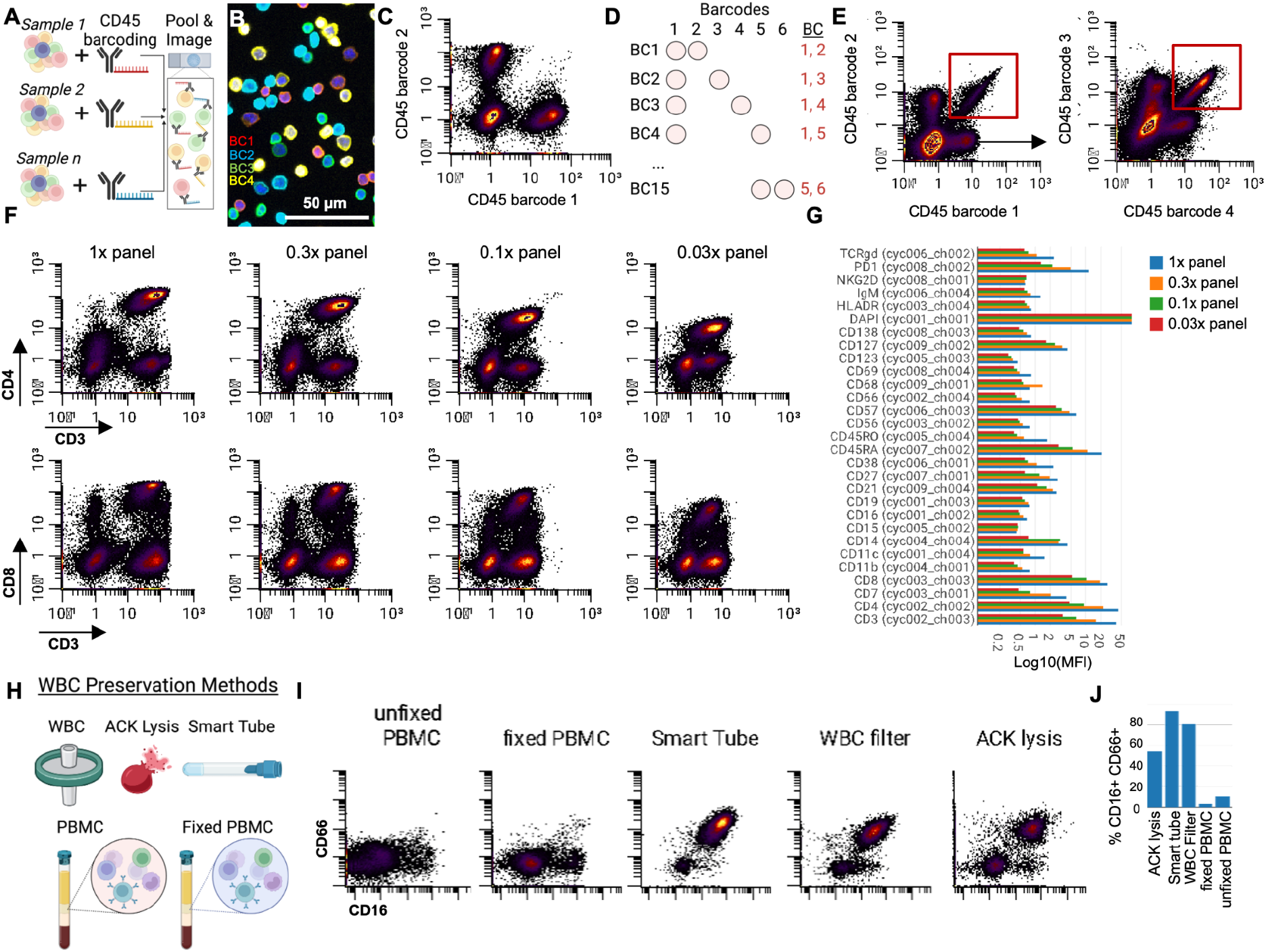
We demonstrate cell and sample-based barcoding for panel titration and different WBC preservation methods. **A)** We barcode cells within different samples by staining with a unique CD45 barcode. **B)** A representative image from four of the barcode channels used in PBMC experiments. **C)** Quantification of barcode fluorescent intensity per cell for two exclusive barcodes. **D)** Combinatorial barcoding enhances the number of barcode possibilities. **E)** Use of combinatorial barcoding showing a “common barcode” gated by barcode 1 and 2, and the “not-gate” used to plot barcode combination 3 and 4. **F)** We used this barcoding strategy to titrate our multiplexed antibody panel concentrations in one experiment. **G)** Quantification of signal allows us to determine the concentration to be used for each experiment. **H)** Different methods we tested preserving WBCs prior to analysis. **I)** CD16 versus CD66 measured within the same barcoded experiment for different PBMC methods and **J)** quantified.

This approach allowed data from the pooled cells to be deconvoluted based on their unique CD45 barcodes (**Fig. 5B**). Cell segmentation of the data revealed that the barcoding created robustly labeled barcode channel positive and negative cells (**Fig. 5C**).

Although single-antibody barcoding is effective and could theoretically generate a unique oligonucleotide barcode for each sample, this method is inefficient as it requires use of one barcode or imaging cycle per condition. To conserve imaging cycles while increasing barcoding capacity, we employed a combinatorial barcoding strategy by combining multiple uniquely barcoded anti-CD45 antibodies. (**Fig. 5D**). Using this strategy with six uniquely oligonucleotide-barcoded anti-CD45 antibodies, we generated 15 unique barcode combinations (**Fig. 5D**). We observed that samples were uniquely identified by double positive staining of corresponding anti-CD45 barcode channels (**Fig. 5E**), demonstrating the effectiveness of our combinatorial barcoding strategy in accurately distinguishing multiple samples within a pooled population.

### Titration of multiplexed imaging antibody panels

In addition to allowing simultaneous analysis of multiple donors, our barcoding approach significantly streamlines the titration of highly multiplexed antibody panels. Titrating each antibody individually across samples can be time-consuming and costly. To address this, we employed our barcoding strategy to titrate our 40-plex antibody panel across four different concentrations in a single experiment. This revealed distinct shifts in the fluorescent signal quantified at the single-cell resolution across markers (**Fig. 5F**). For instance, we observed a marked reduction in CD4 signal as the concentration decreased from 1x to 0.3x, while CD8 showed no significant decrease until concentrations dropped from 0.3x to 0.1x (**Fig. 5F**). By quantifying these shifts across all markers, we were able to determine the optimal antibody concentration for staining PBMCs (**Fig. 5G**). Consequently, combinatorial sample barcoding can be accomplished through CD45 antibody staining and pooling of samples, eliminating batch effects and reducing costs per experiment.

### Sample preparation enables capture of representative immune cell populations from blood

A persistent challenge in single-cell immune analyses is the loss of neutrophils, which are notoriously difficult to maintain during sample processing (*29*). We used our barcoding system to test in a single experiment five different immune cell sample preparation methods from the blood of healthy donors: gradient isolated PBMCs, gradient isolated PBMCs plus fixation, blood processed with white blood cell (WBC) filters, whole blood processed with ACK (Ammonium-Chloride-Potassium) lysing of red blood cells, and whole blood processed with smart tube cell fixation (**Fig. 5H**).

Using our anti-CD45 barcoding approach, we pooled, stained with a panel of 39 antibodies, and imaged all conditions simultaneously. Part of this antibody panel we included two markers that are expressed by human neutrophils: CD16 and CD66. The results showed substantial differences between the sample preparation methods (**Fig. 5I**). Quantification of neutrophils as a percentage of total cells confirmed that the fixed and unfixed PBMC methods retained very few neutrophils, consistent with previous reports (*30, 31*). In contrast, the other methods preserved significantly higher proportions of neutrophils (**Fig. 5J**). These differences extended to other immune cell populations as well (**Fig. S4**). All samples were performed with WBCs that were isolated and stored at −20C. Since neutrophil percentages in healthy human blood range from 40-70%, the ACK lysis preparation strategy of −20C stored samples resulted in value within this range, where smart tube and WBC filters may exaggerate neutrophils abundance (possibly due to fragmentation) with values closer to 80% for the same donor. This data shows that using this approach we can successfully barcode human WBCs and perform quantitative assessments with different WBC preservation methods, including those that retain granulocyte fractions.

## Discussion

While multiplexed imaging techniques have been developed and published to allow reproducibility by other labs (*7–10*), many of these approaches rely on expensive close-box microscopy solutions, limited or almost no automated fluidics capabilities, and lack of open-source flexibility for novel spatial-omics assay developers. These limitations have hindered the broader adoption of these technologies, particularly in resource-limited settings or for field applications, such as during infectious disease outbreaks.

Fluid-Squid addresses these challenges by providing a low-cost, customizable, and open-source platform that integrates automated fluidics with advanced microscopy. Its versatility in imaging both tissues and cells, combined with its portability, offers significant advantages for labs looking to adopt or adapt this system for novel applications. For example, Fluid-Squid’s compatibility with non-slide-based imaging means much larger tissue sections can be analyzed, and cells cultured over extended periods of time can be characterized in situ. Furthermore, our demonstration that CODEX antibody panels can be lyophilized and reconstituted expands the potential for field-based and global applications, where cold-chain storage may not be feasible.

In this study, we also advanced techniques by applying CODEX multiplexed imaging to dissociated single cells, specifically PBMCs and other WBC preservation methods, for the first time. This method offers multiplexing at scale while preserving critical features such as cell and nuclear morphology and the spatial localization of targets within individual cells—advantages that other single-cell approaches often lose.

The integration of cellular barcoding within our system further enhances its utility by enabling the simultaneous profiling of multiple samples. This feature allowed for rapid titration of a 40+ antibody panel in a single experiment. Moreover, it enables reducing both time and costs while minimizing batch effects. This barcoding capability holds particular promise for large-scale cohort studies or longitudinal studies in fields such as immunology, cancer therapy, and infectious disease monitoring – both in lab and field settings. Furthermore, combining this barcoding strategy with emerging pooled perturbation methods, such as Perturb-View (*32*), opens up new possibilities for multiplexed perturbation experiments and large-scale functional studies. Similarly, this method could provide insights into cell-cell interactions and spatially defined ligands such as the immune synapse.

Despite its many advantages, Fluid-Squid currently has some limitations as well. The fluidics system is presently restricted to 12 automated cycles per experiment, which may be insufficient for some highly multiplexed protocols. Efforts are underway to improve the fluidics system to handle more cycles and enhance its robustness across various buffer conditions, broadening its applicability beyond CODEX multiplexed imaging. Additionally, the current microscope setup supports only four-color imaging, but future iterations could incorporate spectral imaging technology to increase the number of channels and expand the system’s multiplexing capacity. Finally, while we have made significant progress in developing image analysis tools and workflows *(10, 18, 19, 22-26, 33)*, there is still room to improve the accessibility and flexibility of these toolsets. As Fluid-Squid continues to evolve, these analysis pipelines will be crucial for the seamless integration of image processing and data interpretation.

In conclusion, Fluid-Squid is an open-source platform that empowers researchers to perform customizable, cost-effective multiplexed imaging of both single cells and tissues. By combining flexibility, scalability, and affordability, Fluid-Squid offers a powerful tool for generating new biological insights across a wide range of research fields. Our approach highlights the role of open-source hardware in biological explorations, paving the path towards an equitable distribution and creation of knowledge.

## Materials and Methods

### Fluid-Squid Setup

#### Squid setup

The Squid microscope used in this work includes motorized XY and Z stages, a Nikon 20x/0.75 Plan Apo objective, a 75 mm machine vision lens that serves as the tube lens, a Sony IMX226 camera, a 850 nm laser autofocus module, Chroma ZT405/488/561/640rpcxt-UF2 dichroic, Chroma ZET405/488/561/640xv2 excitation filter and Chroma ZET405/488/561/640mv2 emission filter, a custom 405/488/530/640 laser engine with SMA fiber connector and a critical illuminator using 400 um square core fiber.

#### Fluidics setup

The Fluidic system used in this work includes a custom PCB, a teensy 4.1 microcontroller, a TTP Ventus disc pump with driver daughter board, honeywell pressure sensors, Sensirion flow sensors, an IDEX 24 port selector valve (HT2425-915-3), clippard solenoid and media isolation valves, and custom machined interfaces for well plates and eppendorf tubes.

#### Software for Fluid-Squid

The python-based software for the first generation fluidics system is available at https://github.com/hongquanli/fluidics-controller-software. The latest microscope software with fluidics system integration can be found at https://github.com/cephla-lab/squid.

### CODEX antibody conjugation and panel creation

#### Antibody conjugation

Each antibody was conjugated to a unique oligonucleotide barcode using our previously established protocol (*10*), after which the tissues were stained with the antibody–oligonucleotide conjugates and we validated that the staining patterns matched the expected patterns already established for immunohistochemistry within positive control tissues of the intestine or tonsil. Otherwise, expected co-expression of markers in the case of staining of suspension cells was taken into account for validating separate cell type and phenotype antibodies. First, antibody–oligonucleotide conjugates were tested in low-plex fluorescence assays and the signal-to-noise ratio was also evaluated at this step, then they were tested all together in a single CODEX multicycle.

#### Lyophilization of CODEX antibodies

An aliquot of the CODEX antibodies for anti-CD8, -B220, -TCRb were taken and rapidly frozen by placing them in liquid nitrogen for 5-10 minutes. Tubes were then transferred to the lyophilizer and allowed to lyophilize until all moisture was removed over 24 hours. Antibodies were then stored at −80 C until used. Then antibodies were reconstituted with double-distilled water to the original constitution with thorough pipetting, spinning down, and allowing 15 minutes before use in staining. The same amount of this aliquot was used to stain one mouse spleen section as was the amount from the original CODEX antibodies that were maintained in the antibody stablizer solution at 4 C.

### Multiplexed Imaging of tissues

#### Tissue sectioning and antibodies

Tissues were individually frozen in OCT molds and then cut on the cryostat and sectioned at a width of 7 μm onto a well of a poly-L-lysine pre-coated 12-well glass-bottom plate. An antibody panel was previously constructed and optimized to image these tissues (*18*). Detailed information about the amount of antibodies used and exposure times is found in Supplementary Table 1.

#### CODEX imaging

The tissue arrays were then stained with the complete validated panel of CODEX antibodies and imaged (*10*). In brief, this entails cyclic stripping, annealing and imaging of fluorescently labeled oligonucleotides complementary to the oligonucleotide on the conjugate. After validation of the antibody–oligonucleotide conjugate panel, a test CODEX multiplexed assay was run, during which the signal-to-noise ratio was again evaluated, and the optimal dilution, exposure time and appropriate imaging cycle was evaluated for each conjugate.

### PBMC/Splenocyte Staining and Multiplexed Imaging

#### PBMC preservation and Thawing

Peripheral blood mononuclear cells (PBMCs) were cryopreserved in media containing 10% DMSO at a density of approximately 10 million cells per mL and stored in liquid nitrogen until use. Six vials of cryopreserved PBMCs (from the same donor) were thawed and processed for staining. PBMC vials were thawed in a 37°C water bath and transferred dropwise to pre-warmed cell thawing medium (RPMI + 10% FBS).

#### Splenocyte staining

Murine cells were obtained from adult mouse lymph nodes and spleens. Obtained cells were treated with ACK lysis buffer to lyse red blood cells, and lysates were filtered through cell strainers to isolate splenocytes. All following steps were performed with the same protocol as the PBMCs (names are exchangeable in the protocol).

#### Cell Preparation and Fixation

Cells were centrifuged at 500g for 5 minutes at room temperature, and the supernatant was carefully decanted. This wash step was repeated twice, first with cell thawing medium, then with Buffer S1. The cells were pooled, centrifuged, and resuspended in Buffer S1. Cells were fixed by resuspending in Buffer S1 containing 1.6% paraformaldehyde (PFA) for 10 minutes at room temperature. The fixed cells were then washed twice with Buffer S1 and prepared for barcoding.

#### Cell Barcoding

The fixed cells were blocked in S2 blocking buffer and split into six tubes for barcoding. Each cell suspension was incubated for 1 hour at room temperature with 1 µL of a unique CD45 barcoding antibody per 100 µL of cells. Following barcoding, the cells were washed three times with Buffer S2 and pooled.

#### Pooling and Antibody Panel Staining

A 10 µL aliquot of the pooled cells was diluted 1:10 in Buffer S2 and counted using a disposable hemocytometer. The total cell count was calculated, and 2.5 million cells were used for subsequent antibody panel staining. The barcoded cells were blocked in S2 blocking buffer for 15 minutes, followed by staining with a 20 µL CODEX antibody panel cocktail (per 2.5 million cells) for 1 hour at room temperature. Cells were washed three times with Buffer S2 after staining. The above procedure was also followed for cells which were not barcoded.

#### Post-Stain Fixation and Plating

Following staining, cells were fixed in Buffer S4 containing 1.6% PFA for 10 minutes and washed three times with PBS. Cells were resuspended in PBS and plated onto poly-L-lysine pre-coated 12-well glass-bottom plates. The plates were incubated at room temperature for 30-60 minutes to allow cells to adhere before a final fixation step with freshly prepared BS3 fixative. Cells were stored in Buffer S4 at 4°C until ready for imaging.

#### Multiplexed Imaging

Stained and fixed cells were imaged using a CODEX-Squid system, with buffer and oligonucleotide preparations carried out according to our prior protocol with CODEX multiplexed imaging with cyclic oligonucleotide.

### WBC preservation methods

Four different methods of blood processing were tested to determine compatibility with CODEX-Octopi staining, focusing on the retention of granulocytes and other leukocytes. Blood samples from a single donor were processed using PBMC isolation, WBC filter processing, ACK lysis, and Smart Tube fixation. Following processing, cells were fixed and stored at 4°C in PBS prior to barcoding and staining.

#### PBMC Isolation

Blood was collected in sodium heparin CPT tubes and centrifuged at 1700 x g for 20 minutes at 21°C with slow acceleration and brake. The upper layer was transferred to 15 mL conical tubes, diluted with PBS, and centrifuged at 300 x g for 15 minutes at 4°C. Cells were washed in PBS, centrifuged, and fixed in 1.6% paraformaldehyde (PFA) for 10 minutes at room temperature. Fixed cells were washed, resuspended in PBS, and divided into aliquots. One aliquot was stored at 4°C, and the other was frozen at −20°C.

#### WBC Filter Processing

Blood was collected in EDTA vacutainers and passed through an Acrodisc WBC filter by gravity. The filter was washed with PBS, and WBCs were eluted by pressing PBS through the filter into a 15 mL conical tube. Cells were centrifuged at 300 x g for 10 minutes at 4°C, fixed in 1.6% PFA for 10 minutes at room temperature, and washed. The cell pellet was resuspended in PBS, divided into two aliquots, with one stored at 4°C and the other frozen at −20°C.

#### ACK Lysis Processing

Blood collected in EDTA vacutainers was mixed with ACK lysis buffer and incubated at room temperature for 3-5 minutes to lyse red blood cells. Cells were centrifuged at 300 x g for 15 minutes at 4°C, washed with PBS, and centrifuged again. The cell pellet was fixed in 1.6% PFA for 10 minutes at room temperature, washed, and resuspended in PBS. Aliquots were stored at 4°C and −20°C.

#### Smart Tube Fixation

Blood collected in sodium heparin vacutainers was mixed with Smart Tube Proteomic Stabilizer in cryovials and incubated for 10 minutes at room temperature. Cryovials were then frozen at −80°C. For thawing, samples were incubated in Thaw-Lyse buffer for 15 minutes at room temperature, centrifuged at 600 x g for 5 minutes at 22°C, and washed twice with Thaw-Lyse buffer. Cells were resuspended in PBS, divided into aliquots, and stored at 4°C and −0°C.

### Quantification of single-cell imaging datasets

#### Fluid-Squid Image Processing

For regular CODEX multiplexed imaging using the Akoya Phenocycler imager, we used the image processing of stitching, image registration, cycle concatenation, and background subtraction built into the commercial software.

#### Cell Segmentation

Fo CODEX multiplexed imaging data from the Akoya Phenocycler imager we used CellVisionSegmenter, a neural network R-CNN-based single-cell segmentation algorithm (*34*). The only parameter that was altered was the growth pixels of the nuclear mask, which we found experimentally to work best at a value of 3.

#### Debarcoding and Gating

We used cellengine.com platform to identify the barcoded populations of cells. In cases where we used one barcode, we would create gates for each marker positivity and exclude these cells from other cell barcodes. For double barcodes we selected only double-positive cells. In experiments we identified cell types based on binary expression of known marker expression in a similar way to the way we do this for flow cytometry or CyTOF.

#### Unsupervised Clustering

Nucleated cells were selected by gating Hoechst 1 and Hoechst 2 double-positive cells (across multiple cycles), followed by z-normalization of protein markers used for clustering (some phenotypic markers were not used in the unsupervised clustering). Cells positive (z > 1) for greater than 35 fluorescent markers were removed from the data. Then the data were overclustered Leiden-based clustering with the scanpy Python package (v.1.9.1). These processing steps were performed based on an in-depth analysis of normalization techniques and unsupervised clustering algorithms used for CODEX multiplexed imaging data of the intestine (*24*). Clusters were assigned a cell type on the basis of average cluster protein expression and the location within the image. Impure clusters were split or reclustered after mapping back to the original fluorescence images.

## Author contributions

J.W.H., H.L., D.R.M., G.P.N., and M.P. Conceptualized, designed, and created the system and designed experiments. J.W.H., H.L., C.C., H.T.L, J.S.F., E.T., B.D-K. and D.R.M., performed experiments and collected data. H.L. designed and built the microscope, H.L., K.M., Y.Y., P.M. developed and iterated the fluidics systems, H.L., K.M. developed the image processing code, J.W.H., H.L., D.R.M., K.M., and C.C. analyzed data. J.W.H., H.L., D.R.M., G.P.N., and M.P. wrote and revised the manuscript.

## Data and materials availability

All data are available upon request by authors.

## Supplementary Materials

Figs. S1 to S5 are provided in the Supplemental Materials document.

