## Supplemental Information for "Fluid-Squid: DIY Multiplexed Imaging of Cells and Tissues"

John W. Hickey *et al.*

**This PDF file includes:** Figs. S1 to S5

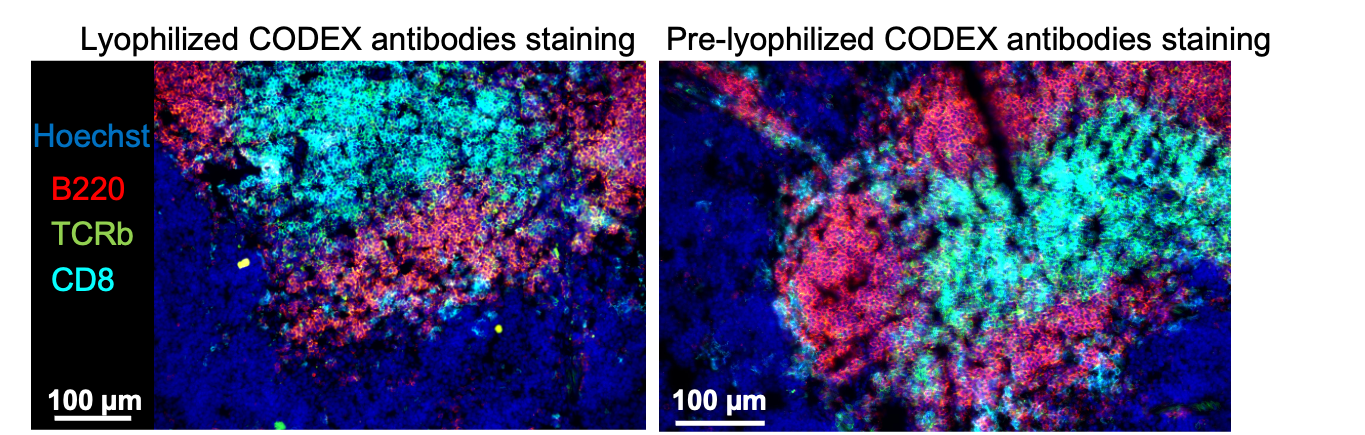

**Fig. S1**: CODEX multiplexed imaging of a mouse spleen using three antibodies that were either kept in solution or lyophilized and resuspended.

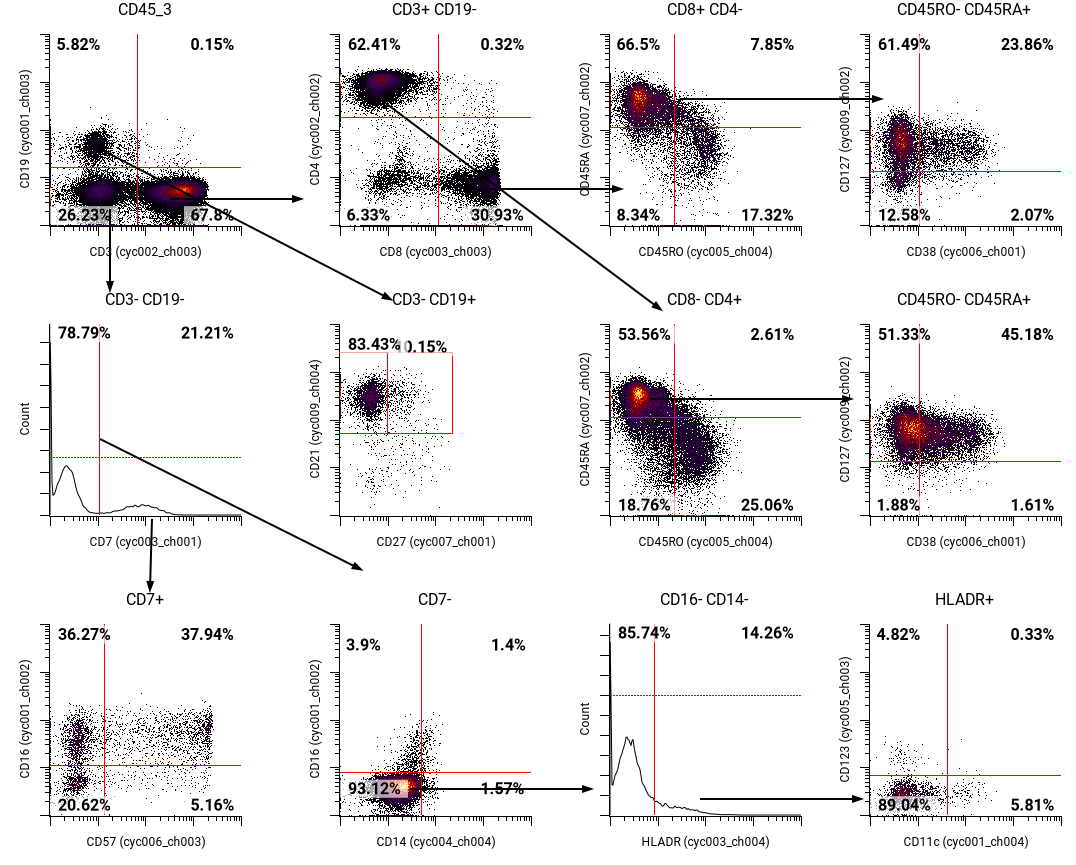

**Fig. S2**: Gating strategy for the CODEX multiplexed imaging of suspension cell experiment (Figure 3).

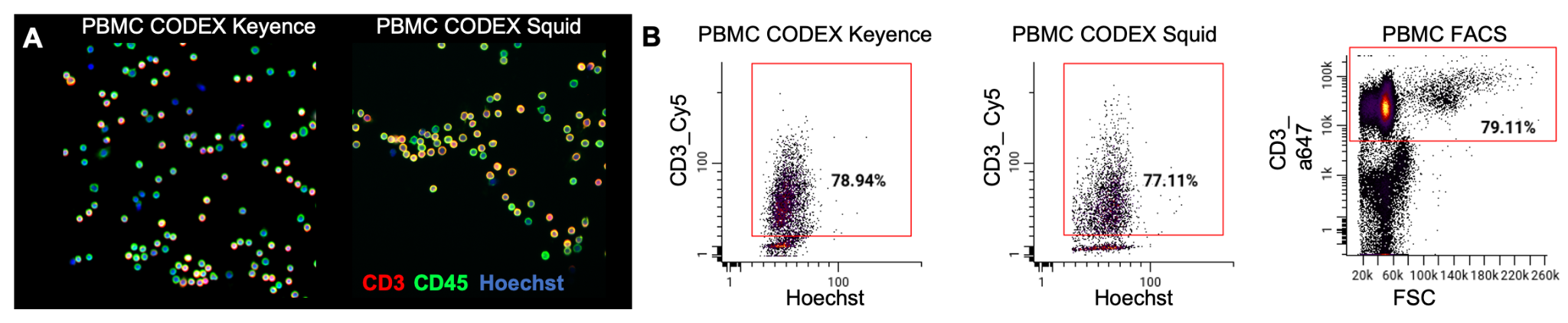

**Fig. S3:** Squid imaging of CODEX multiplexed PBMC staining compared to a Keyence microscope. A) Images and B) quantification of gated CD3+ cells assessed by CODEX Keyence, CODEX Squid, and flow cytometry.

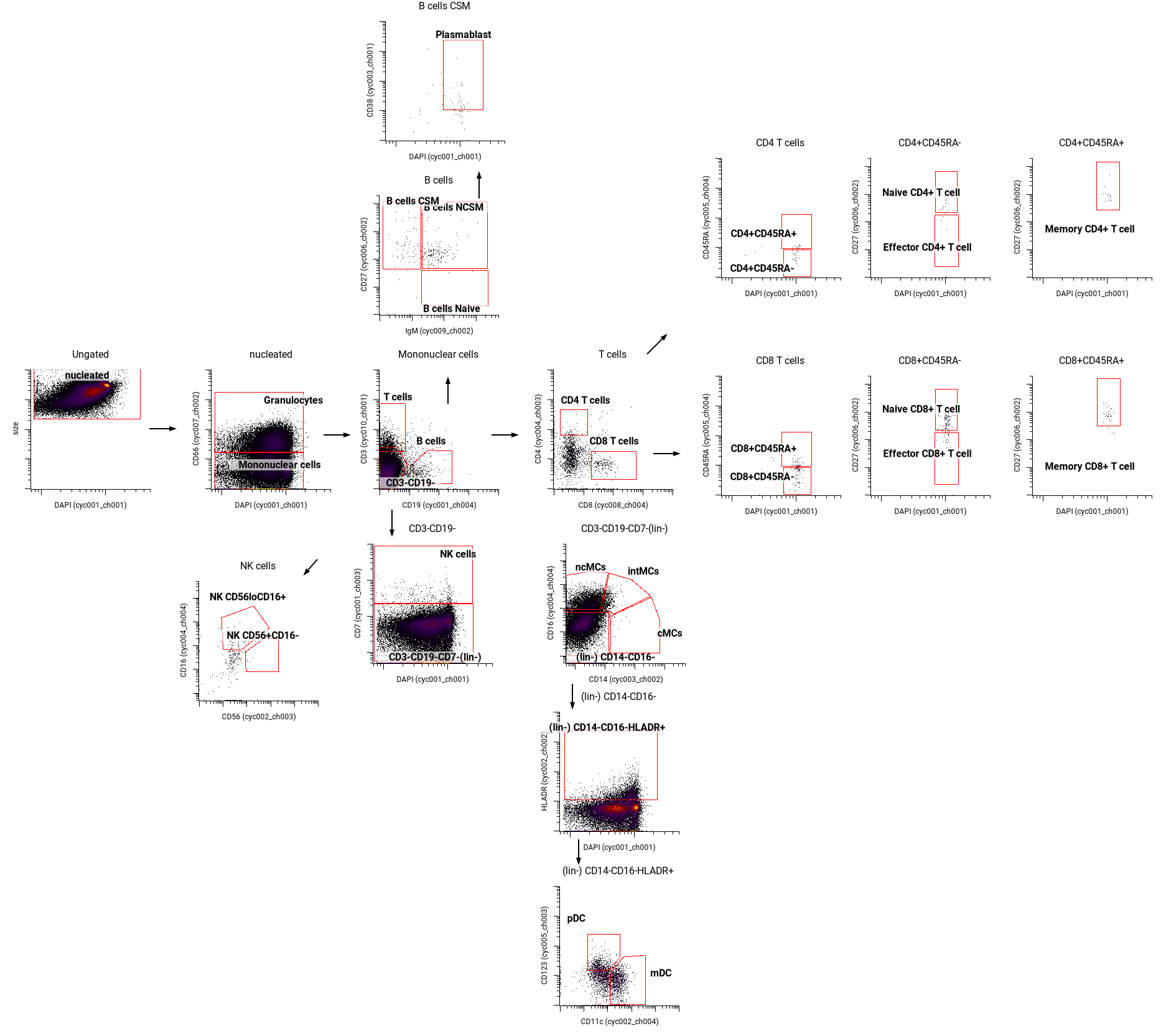

**Fig. S4:** Gating strategy for characterizing the different blood preparation strategies post cell debarcoding.

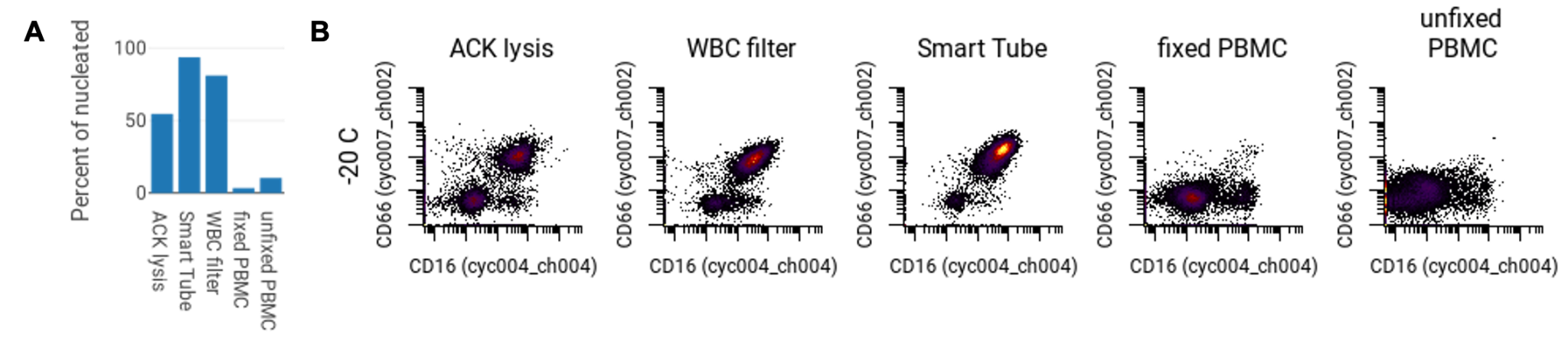

**Fig. S5:** Preparation and storage temperatures for characterizing WBCs **A)** Percent of CD66+, CD16+ cells of all cells imaged. **B)** Plots of CD66 by CD16 for different methods and storage temperatures used.
